# Maturation of Purkinje cell firing properties relies on granule cell neurogenesis

**DOI:** 10.1101/2020.05.20.106732

**Authors:** Meike E. van der Heijden, Elizabeth P. Lackey, Fatma S. Işleyen, Amanda M. Brown, Ross Perez, Tao Lin, Huda Y. Zoghbi, Roy V. Sillitoe

**Affiliations:** Department of Pathology & Immunology, Baylor College of Medicine, Houston, Texas, USA; Department of Neuroscience, Baylor College of Medicine, Houston, Texas, USA; Program in Developmental Biology, Baylor College of Medicine, Houston, Texas, USA; Development, Disease Models & Therapeutics Graduate Program, Baylor College of Medicine, Houston, Texas, USA; Jan and Dan Duncan Neurological Research Institute at Texas Children’s Hospital, Houston, Texas, 77030, USA; University of St. Thomas, Houston, Texas, USA; Howard Hughes Medical Institute, Department of Molecular and Human Genetics, Baylor College of Medicine, Houston, Texas, USA

**Keywords:** Purkinje cells, granule cells, cerebellum, electrophysiology, development, behavior, cerebellar circuit, *Atoh1*, cerebellar function

## Abstract

Preterm infants that suffer cerebellar insults often develop motor disorders and cognitive difficulty. Granule cells are especially vulnerable, and they likely instigate disease by impairing the function of Purkinje cells. Here, we use regional genetic manipulations and *in vivo* electrophysiology to test whether granule cells help establish the firing properties of Purkinje cells during postnatal mouse development. We generated mice that lack granule cell neurogenesis and tracked the structural and functional consequences on Purkinje cells in these agranular pups. We reveal that Purkinje cells fail to acquire their typical connectivity and morphology, and the formation of characteristic Purkinje cell firing patterns is delayed by one week. We also show that the agranular pups have impaired motor behaviors and vocal skills. These data argue that granule cell neurogenesis sets the maturation time window for Purkinje cell function and refines cerebellar-dependent behaviors.

## INTRODUCTION

Abnormal cerebellar development instigates motor diseases and neurodevelopmental disorders including ataxia, dystonia, tremor, and autism. These conditions are highly prevalent in premature infants and in newborns with cerebellar hemorrhage (Dijkshoorn et al., 2020; Limperopoulos et al., 2007; Steggerda et al., 2009; Zayek et al., 2012), who ultimately attain a smaller cerebellar size compared to children born full-term (Limperopoulos et al., 2005; Volpe, 2009). During the third trimester of human development, which corresponds to the first two postnatal weeks in mice (Sathyanesan et al., 2019), the cerebellum increases five-fold in size due to the rapid proliferation of granule cell precursors and the integration of granule cells into the cerebellar circuit (Chang et al., 2000). Observations from clinical data indicate a strong correlation between cerebellar size and cognitive disorders, suggesting that this period of cerebellar expansion is a critical developmental time-window for establishing cerebellar function. Pig and mouse studies confirm that preterm birth, postnatal hemorrhage and hypoxia all result in lower granule cell numbers (Iskusnykh et al., 2018; Yoo et al., 2014) and abnormal motor control (Sathyanesan et al., 2018; Yoo et al., 2014). Importantly, such peri- and postnatal insults are accompanied by impairments in the intrinsic firing properties of Purkinje cells (Sathyanesan et al., 2018). Purkinje cells are the sole output of the cerebellar cortex and integrate input from up to two hundred fifty thousand excitatory granule cell synapses (Huang et al., 2014), though the predominant Purkinje cell action potential called the simple spike, is intrinsically generated (Raman and Bean, 1999). In this context, it is intriguing that genetically silencing granule cells caused modest alterations to the baseline firing properties of Purkinje cells and impaired only the finer aspects of motor learning, but not gross motor control (Galliano et al., 2013). The discordance between the phenotypes in mutant mice with lower granule cell numbers and mice lacking granule cell function questions whether granule cell neurogenesis, rather than granule cell synaptic signaling, drives the maturation of Purkinje cell firing *in vivo*.

To probe these cellular interactions, we manipulated the mouse cerebellum by genetically blocking granule cell neurogenesis. We used the *En1* lineage to delete the proneural gene, *Atoh1*, from the hindbrain. *Atoh1* is necessary for granule cell development (Ben-Arie et al., 1997) but it is not expressed in Purkinje cells. In this agranular model, we test how Purkinje cells develop their anatomy and function using immunohistochemistry and *in vivo* electrophysiology recordings in the second postnatal week. We further investigated the motor and vocal skills of the agranular pups to test how the structural and functional changes impact the expression of normal behaviors.

## RESULTS

### Mice lacking *Atoh1* from the *En1* domain do not form differentiated granule cells

To test the hypothesis that granule cell neurogenesis is essential for the functional development of Purkinje cells, we first established a model of agranular mice that is not initiated by the cell-autonomous development of Purkinje cells. In previous models, agranular mice lack granule cells due to spontaneously occurring mutations in genes with widespread expression patterns. In those mice, therefore, one cannot differentiate cell-extrinsic from cell-intrinsic effects (Dusart et al., 2006; Gold et al., 2007). Instead, we made use of the distinct origins of Purkinje cells and granule cells. (**Figure 1A-B**) (Hoshino et al., 2005; Rose et al., 2009). The intersection of the *Atoh1* and *En1* lineages converges on granule cells (Wang et al., 2005), but not Purkinje cells (**Figure 1C**). *Atoh1* is necessary for the development of granule cells, the most populous cell type in the cerebellum, but *Atoh1* null mice are neonatal lethal (Ben-Arie et al., 1997). *En1* is a homeobox transcription factor that is expressed in the mesencephalon and rhombomere 1 by embryonic day E(8) where it is required for the formation of the cerebellum (Davis and Joyner, 1988; Wurst et al., 1994). Considering their spatial and temporal expression domains, it was previously published that conditional deletion of *Atoh1* from the *En1* domain produces viable mice (*En1^Cre/+^;Atoh1^fl/-^*) with a remarkably small cerebellum (van der Heijden and Zoghbi, 2018). We confirm these findings (**Figure 1D**) and show that postnatal *En1^Cre/+^;Atoh1^fl/-^* mice lack a densely packed granule cell layer (**Figure 1E**). Calbindin staining shows that *En1^Cre/+^;Atoh1^fl/-^* mice do have Purkinje cells, although they do not settle into a monolayer, as is characteristic of the normal cerebellum. At the gross anatomy level, the mediolateral morphological cerebellar divisions and normally obvious lobules and deep fissures in the anteroposterior axis fail to form (**Figure 1F-H**). Interestingly, we observed *En1^Cre/+^;Atoh1^fl/-^* Purkinje cells that were ectopically located rostral to the cerebellum, in the inferior colliculus (from here on, referred to as displaced Purkinje cells), suggesting abnormal settlement due to over-migration. Analyses of molecular markers expressed by different cerebellar neuron subtypes confirm that the deletion of *Atoh1* depletes differentiated granule cells (**Supp. 1A**). Consistent with the overall smaller size of cerebellum, the mutation impacts the pool of excitatory unipolar brush cells that are localized to lobule IX and X in control mice, interneurons that are also derived from the *Atoh1* lineage (**Supp. 1B-D**). The representation of different inhibitory neurons, including Purkinje cells, remains robust and clearly detected by cell-type specific markers (**Supp. 1E-H**). These data indicate that although there is an equivalent reduction in cerebellar morphology and cytoarchitecture in the mutant, the principle defect is the elimination of granule cells. We conclude that *En1^Cre/+^;Atoh1^fl/-^* mutant mice are a unique model for cerebellar agranularity because the primary defect is independent of genes expressed in the Purkinje cells.

**Figure 1.**
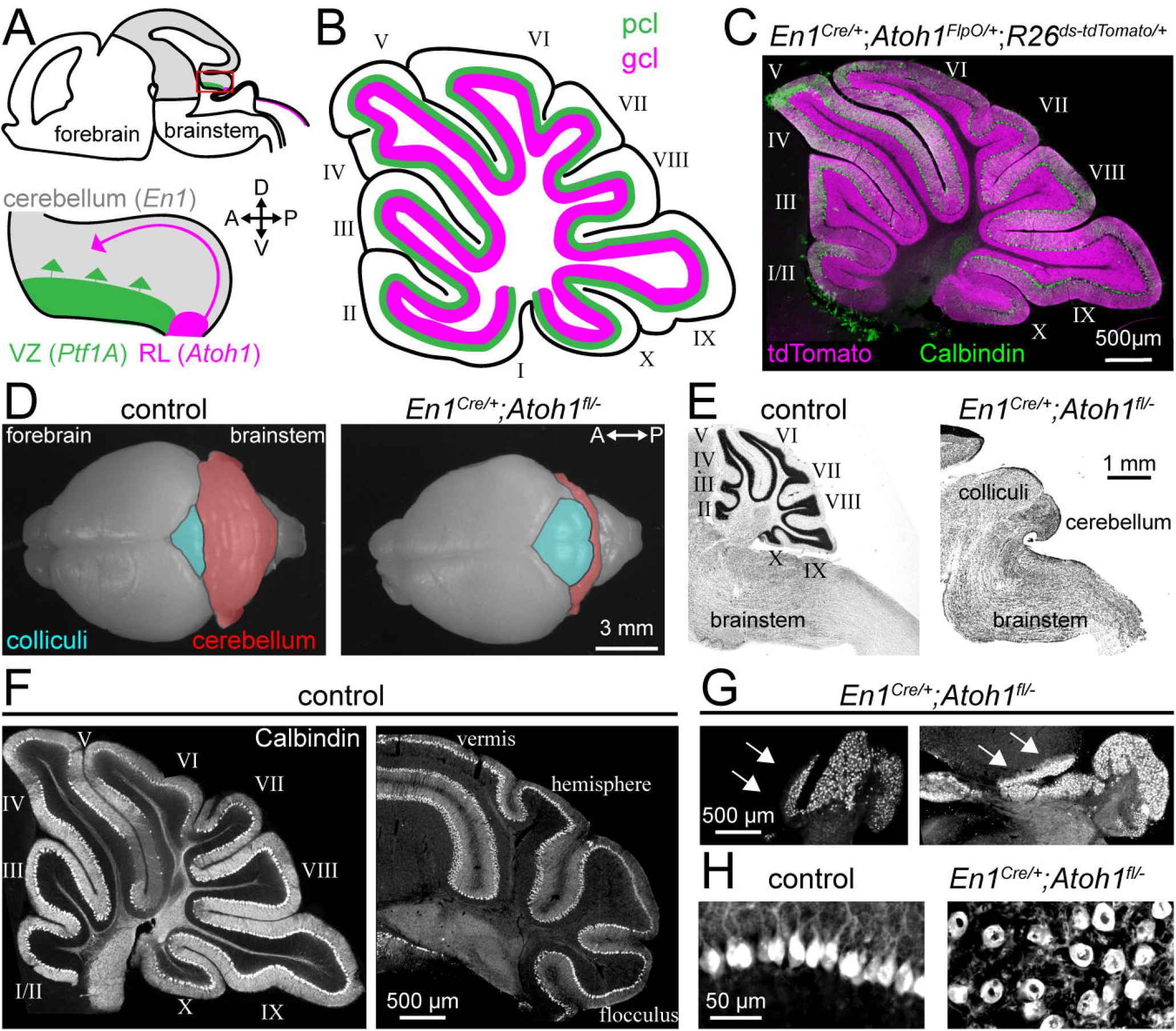
Conditional deletion of *Atoh1* from the *En1* domain results in an agranular cerebellum. **A.** Schematic of an embryonic brain. Inset is the cerebellar anlage. *Atoh1* domain (granule cell precursors, pink), *Ptf1a* domain (Purkinje cell precursors, green), *En1* domain (grey). Orientation is the same for all panels unless otherwise indicated. **B.** Schematic of a sagittal section of a P14 cerebellum. Purkinje cell=green; granule cell=pink. Cerebellar lobules are labelled with Roman numerals (Larsell, 1952). **C.** Intersectional labelling of *En1;Atoh1* domain with tdTomato (pink) shows no overlap with Purkinje cells (Calbindin; green). **D.** Whole brain images of control and *En1^Cre/+^;Atoh1^fl/-^* mice showing abnormal gross morphology. **E.** Sagittal sections of control and *En1^Cre/+^;Atoh1^fl/-^* hindbrains stained with cresyl violet to visualize cell nuclei. **F.** and **G.** Sagittal and coronal sections of P14 cerebella of control (**F**) and *En1^Cre/+^;Atoh1^fl/-^* (**G**) mice stained with Calbindin (grey). Arrows indicate Purkinje cells that have migrated into the colliculi. **F.** and **G.** are presented at the same magnification. **H.** Higher magnification images of Calbindin staining in control and *En1^Cre/+^;Atoh1^fl/-^* mice. Images are representative for N=3 brains for each genotype.

### Purkinje cells in *En1^Cre/+^;Atoh1^fl/-^* mice display hallmarks of anatomical immaturity

Next, we set out to investigate whether Purkinje cells in our *En1^Cre/+^;Atoh1^fl/-^* model had abnormal excitatory inputs and morphology (summarized in **Figure 2A**). Early postnatal Purkinje cells receive direct synaptic contacts from mossy fibers originating from various nuclei in the brainstem and spinal cord (Sillitoe, 2016) and from multiple climbing fibers originating in the inferior olive, although the direct innervation from granule cells has yet to form parallel fibers (Lackey et al., 2018; Mason and Gregory, 1984; White and Sillitoe, 2013). After circuit reorganization, mossy fibers no longer contact Purkinje cells, only one climbing fiber innervates each Purkinje cell, and thousands of inputs from parallel fibers now dominate the Purkinje cell dendrite. We anatomically examined synapses in postnatal day (P)14 Purkinje cells, as most synaptic rearrangements occur before this timepoint. We found that Purkinje cells in *En1^Cre/+^;Atoh1^fl/-^* mice have a significant reduction in Vglut1-positive inputs in both cerebellar and displaced Purkinje cells (**Figure 2B**; parallel fibers/mossy fibers). Conversely, there was dense staining for Vglut2-positive inputs to cerebellar and displaced Purkinje cells (**Figure 2C**; climbing fibers/mossy fibers). To test if some inputs were from the spinal cord that sends an early major cerebellar projection, we injected the anterograde tracer WGA-Alexa 555 (Gebre et al., 2012; Lackey and Sillitoe, 2020) into the lower thoracic-upper lumbar spinal cord of P12 mice and observed WGA-Alexa 555 labeled fibers and terminals at P14 after two days of tracer transport. In control cerebella, mossy fibers project to the cerebellar cortex in a striped pattern that respects the topography of ZebrinII, a molecular marker of medial-lateral Purkinje cell patterns (**Figure 2D, E** and **Supp. 2A-G**) (Brochu et al., 1990; Sillitoe and Hawkes, 2002). In the *En1^Cre/+^;Atoh1^fl/-^* cerebellum, ZebrinII is organized in clusters rather than sharp stripes, in a pattern that resembles the normal early neonatal architecture (**Supp. 2H-N**) (Fujita et al., 2012; Sugihara and Fujita, 2013). Interestingly, we found that spinocerebellar mossy fibers projected mainly to ZebrinII-negative domains in the *En1^Cre/+^;Atoh1^fl/-^* mice. The labeled mossy fibers were associated with Purkinje cells of the same ZebrinII identity irrespective of whether they were located within the cerebellum or ectopic and displaced in the colliculi (**Figure 2E**), which is similar to their association with ZebrinII-negative stripes in control animals.

**Figure 2.**
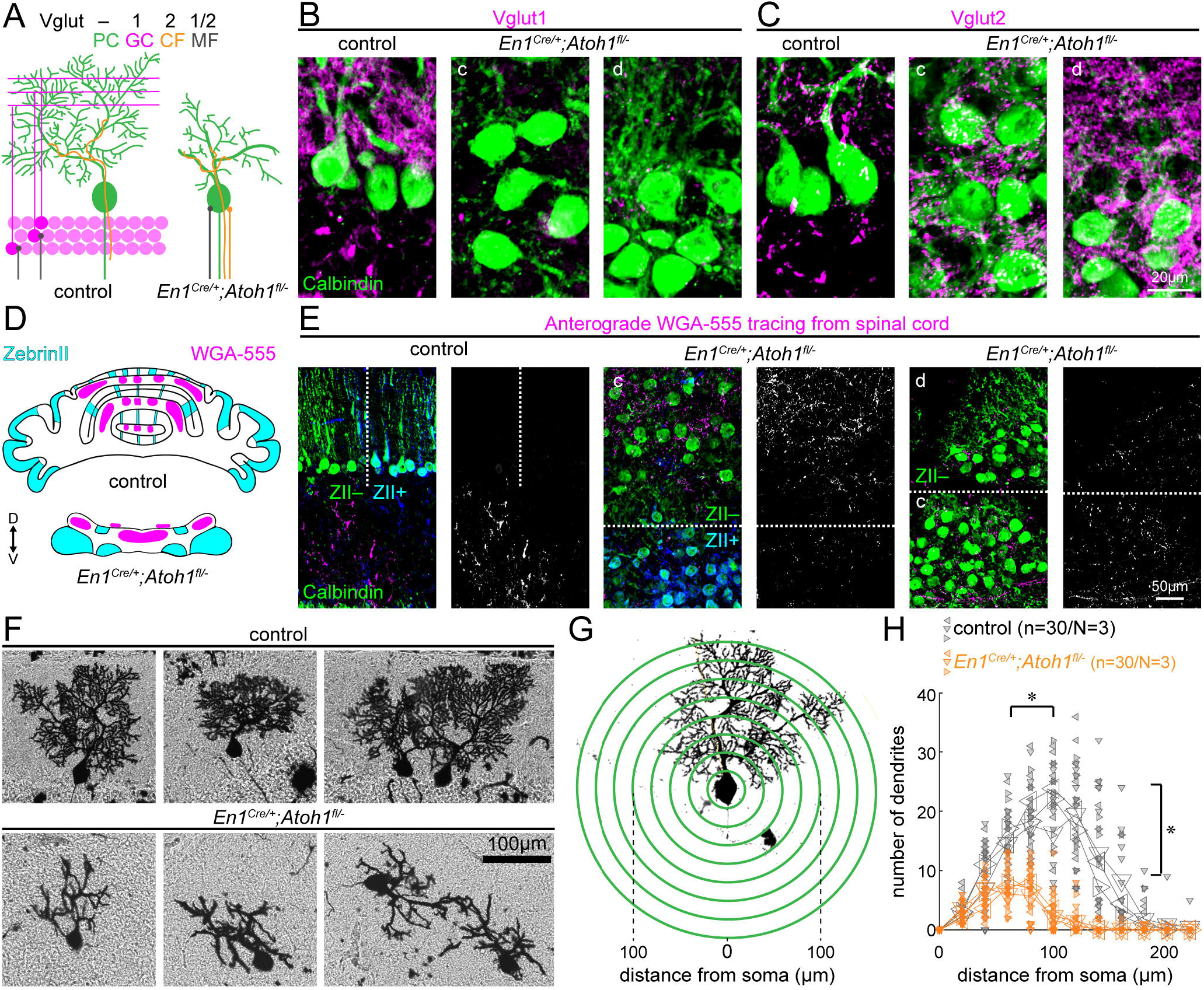
Abnormal glutamatergic input and morphology of *En1^Cre/+^;Atoh1^fl/-^* Purkinje cells. **A.** Schematic of a control Purkinje cell and an *En1^Cre/+^;Atoh1^fl/-^* Purkinje cell (based on results from **B-H**). MF=Mossy Fiber; CF=Climbing fiber. **B.** Images of Calbindin (green) and Vglut1 (pink) staining. For *En1^Cre/+^;Atoh1^fl/-^* mice in **B, C,** and **E** c=cerebellum, and d=displaced Purkinje cells. Images are representative for N=3 brains for each genotype. **C.** Images of Calbindin (green) and Vglut2 (pink) staining. **D.** Schematic of a control Purkinje cell, the *En1^Cre/+^;Atoh1^fl/-^* ZebrinII staining pattern (see **Supp. Figure 2**) and WGA-Alexa 555 tracing form the spinal cord. **E.** Representative images of WGA-Alexa 555+ terminals in the cerebellum. Dotted lines represent the border between ZebrinII-positive (cyan) and -negative region (left four images) or between the cerebellum and colliculi (right images). Black and white image shows the pattern of WGA-Alexa 555 positive terminals. **F.** Representative images of Golgi-Cox-labelled Purkinje cells in control (top row) and *En1^Cre/+^;Atoh1^fl/-^* brains (bottom row). **G.** Sholl analysis for dendritic complexity. **H.** Purkinje cells in *En1^Cre/+^;Atoh1^fl/-^* mice have shorter and less branched Purkinje cell dendrites (n=30/N=3 for each genotype, each animal is indicated with a differentially oriented triangle). Linear mixed model with genotype as the fixed effect and mouse number as the random effect. *P<0.001 for both distance from soma and branch number. All images were acquired from the cerebellum of P14 mice.

We also examined the morphology of P14 Purkinje cells since previous studies suggest that decreased excitatory input alters Purkinje cell dendrite outgrowth (Bradley and Berry, 1976; Park et al., 2019). Using Golgi-Cox staining, we found that Purkinje cells in *En1^Cre/+^;Atoh1^fl/-^* mice had stunted and smaller dendritic arbors compared to controls (**Figure 2F**). Neighboring Purkinje cells did not orient their arbors in the same direction, as is observed in control cerebella (**Figure 2F**). Sholl analysis revealed that the Purkinje cells in the mutant are smaller with less bifurcated dendritic branches (**Figure 2G** and **H**). In summary, Purkinje cells in *En1^Cre/+^;Atoh1^fl/-^* mice have less morphological complexity and Vglut1-positive synapses, although they do receive Vglut2-positive synapses, some of which are mossy fiber inputs. These anatomical data suggested to us that Purkinje cells in the agranular *En1^Cre/+^;Atoh1^fl/-^* mice may be trapped at an immature stage.

### Lack of granule cells in *En1^Cre/+^;Atoh1^fl/-^* mice blocks the maturation of Purkinje cell firing

Purkinje cells have a distinct firing profile characterized by intrinsically generated simple spikes and climbing fiber-induced complex spikes. *In vitro* recordings in rats showed that Purkinje cell firing properties change significantly during early postnatal development, but it is unclear how firing evolves *in vivo*, during this dynamic period of rewiring (McKay and Turner, 2005). We set out to answer two questions. First, we wanted to know how Purkinje cell firing changes during the dynamic period, and second, whether Purkinje cell firing is affected in *En1^Cre/+^;Atoh1^fl/-^* mice that are deficient in synaptic rewiring. We performed extracellular recordings in anesthetized mice and observed dramatic differences between P14 control and *En1^Cre/+^;Atoh1^fl/-^* mice (**Figure 3A-F**). Purkinje cells in P14 controls fired in burst-like patterns, resulting in a large disparity between the calculated firing frequency and frequency mode (or preferred frequency) (**Figure 3B** and **E**). In contrast, Purkinje cells in *En1^Cre/+^;Atoh1^fl/-^* mice fired relatively regularly, yet at a lower frequency (**Figure 3D** and **F**). We tested the firing features with five parameters: frequency (spikes/recording time; **3G**), frequency mode (most frequently observed frequency; **3H**), CV (a measure for global regularity; **3I**), CV2 (a measure for local, or intrinsic regularity; **3J**), and pause percentage (defined as a discrete portion of the trace during which no spikes were observed; **3K**). We calculated these parameters for Purkinje cells recorded from P7-14 controls and P14 *En1^Cre/+^;Atoh1^fl/-^* mutants and found that frequency and frequency mode gradually increased with age in controls, but that Purkinje cells in the P14 *En1^Cre/+^;Atoh1^fl/-^* mice fired slower than control cells at P10-14 (**Figure 3G-H**). Next, we detected a gradual increase in CV but a decrease in CV2 in control cells. In other words, older cells fired in bursts, although the inter spike interval (ISI) during the bursts became more regular. This trend was not observed in *En1^Cre/+^;Atoh1^fl/-^* mice, with their Purkinje cells having a lower CV than control P12-14 Purkinje cells and higher CV2 than control P8-14 Purkinje cells (**Figure 3I-J**). Additionally, as a result of their increase in burstiness, control Purkinje cells increase the pause proportion with age, a feature that was not observed in the P14 *En1^Cre/+^;Atoh1^fl/-^* mice (statistically different from control P12-14 Purkinje cells) (**Figure 3K**). Finally, we performed a cluster analysis on the first three principle components of the group means for each of the parameters (**Figure 3G-K**). This analysis revealed that Purkinje cells in *En1^Cre/+^;Atoh1^fl/-^* mice have the lowest dissimilarity with control P8 Purkinje cells and that control P7-P10 Purkinje cells form a distinct cluster from their counterparts at P11-14 (**Figure 3L**). Please refer to **Figure Supp. 3** for representative recordings from control P7-14 and *En1^Cre/+^;Atoh1^fl/-^* P14 Purkinje cells.

**Figure 3.**
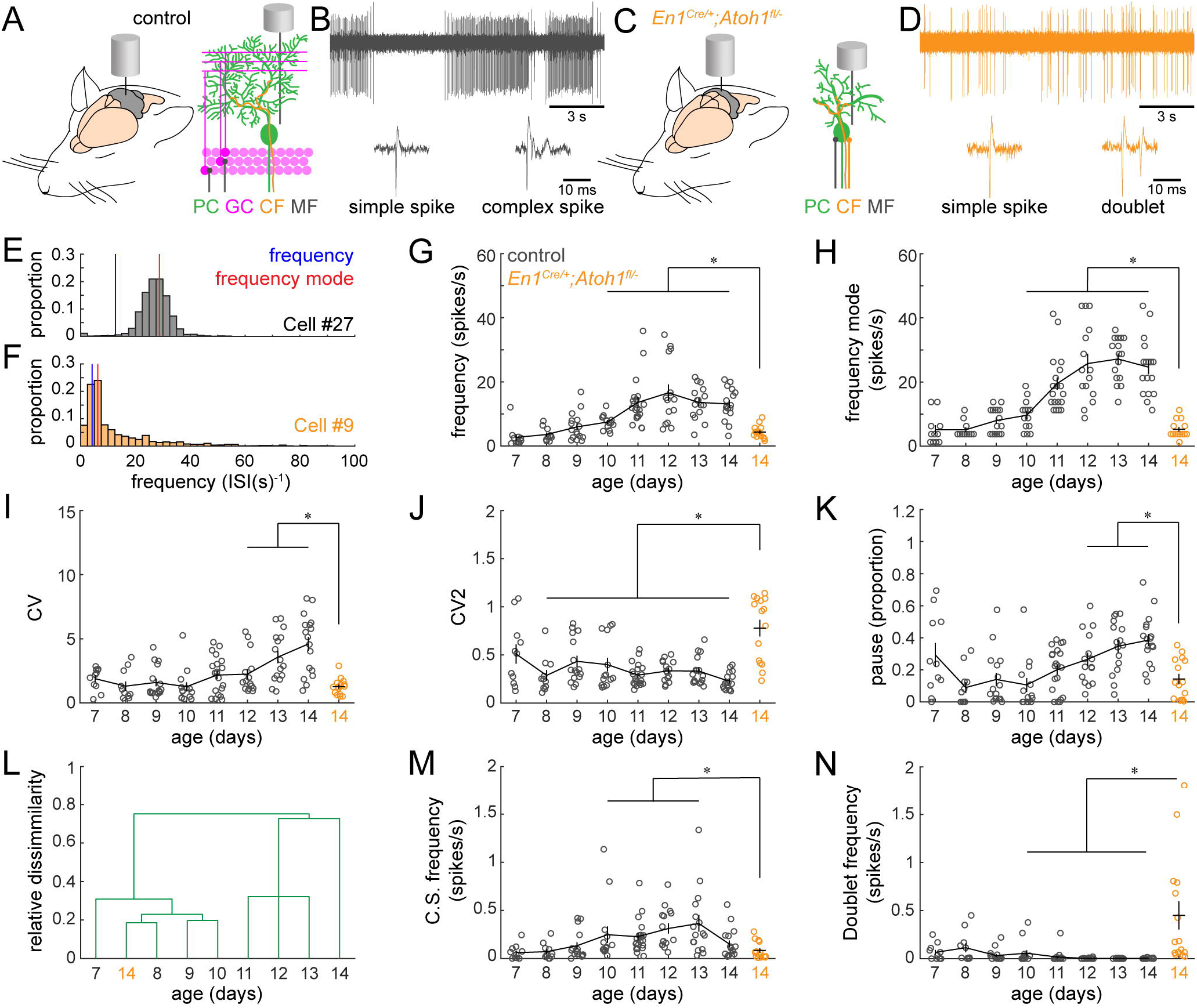
Dynamics of *in vivo* firing properties of Purkinje cells recorded from the cerebellum of control and *En1^Cre/+^;Atoh1^fl/-^* mice. **A.** Schematic of Purkinje cell recordings in control mice. MF=Mossy Fiber; CF=Climbing fiber. **B.** Representative, 15 seconds (s) recording trace from a control P14 Purkinje cell (top). Simple spike and complex spike (bottom). **C.** Schematic of Purkinje cell recordings in *En1^Cre/+^;Atoh1^fl/-^* mice. **D.** Representative, 15 s recording trace from an *En1^Cre/+^;Atoh1^fl/-^* P14 Purkinje cell (top). Simple spike and doublet (bottom). **E.** and **F.** ISI distributions of control (**E**) and *En1^Cre/+^;Atoh1^fl/-^* (**F**) P14 Purkinje cells. Frequency calculated as spikes/s indicated in blue, and frequency mode indicated in red. Cells in **B, D, E,** and **F** were chosen because they most closely represented group averages. **G.** Simple spike firing frequency (spikes/recording time). **H.** Simple spike frequency mode (peak ISI^-1^ distribution). **I.** Simple spike CV (global firing irregularity). **J.** Simple spike CV2 (local firing irregularity). **K.** Pause percentage (proportion of recording with ISI > five times average ISI). **L.** Cluster analysis on the first three Principle Components (accounting for >85% of variation) of the average of intrinsic firing properties (from **G-K**) calculated per age and genotype. **M.** Complex spike firing frequency (spikes/recording time). **N.** Doublet firing frequency (spikes/recording time). For **G-K** and **M-N**, significance was determined using a t-test between Purkinje cells in P14 *En1^Cre/+^;Atoh1^fl/-^* mice and each of the timepoints of Purkinje cells from the control mice. Significance was accepted at P<0.00065 (=0.05/8, for 8 repeated tests). N-numbers: P7: n=11/N=6; P8: n=11/N6; P9: n=16/N=5; P10: n=14/N=7; P11: n=20/N=7; P12: n=15/N=4; P13: n=16/N=4; P14: n=17/N=7; *En1^Cre/+^;Atoh1^fl/-^* P14: n=15/N=6.

We also observed that the climbing fiber-induced Purkinje cell complex spike activity was altered when comparing between control and *En1^Cre/+^;Atoh1^fl/-^* cells. First, the number of classical complex spikes, which are defined by a large sodium spike followed by a train of 3-5 smaller calcium-mediated spikelets (Davie et al., 2008; Zagha et al., 2008), was lower in Purkinje cells of *En1^Cre/+^;Atoh1^fl/-^* mutants (the difference is statistically significant when the mutant is compared to P10-13 controls, **Figure 3M**). Second, the Purkinje cells in *En1^Cre/+^;Atoh1^fl/-^* mice fired distinct “doublets”, which are characterized by an initial simple spike-like action potential, followed by a smaller action potential that occurs within 20 ms. A similar profile of doublets was previously reported in neonatal rats (Puro and Woodward, 1977; Sokoloff et al., 2015). While we observed doublets in both genotypes and all ages studied, they were most frequent in the P14 *En1^Cre/+^;Atoh1^fl/-^* cerebellum (the difference is statistically significant when compared to P10-P14 controls, **Figure 3N**). We postulate that Purkinje cell physiology changes substantially during the period of synaptic rewiring, though many of these changes do not occur in *En1^Cre/+^;Atoh1^fl/-^* mice.

### Circuit defects in *En1^Cre/+^;Atoh1^fl/-^* mice reflect impaired cerebellar-dependent behaviors

The cerebellum controls motor coordination and balance as well as social behaviors including ultrasonic vocalization (USV) in neonatal pups (Fujita et al., 2008; Lalonde and Strazielle, 2015). Interestingly, the contribution of the cerebellum to these behaviors is established before circuit rewiring is completed. Therefore, we were curious to know whether the immature circuit of *En1^Cre/+^;Atoh1^fl/-^* mice was sufficient to perform a substantial repertoire of cerebellar-dependent behaviors. Observations of control and *En1^Cre/+^;Atoh1^fl/-^* mutant mice showed overt phenotypic differences in motor control (**Supp Video 1** and **Figure 4A**). At P14, control mice explore an open arena with smooth intentional motions, whereas *En1^Cre/+^;Atoh1^fl/-^* mice often fall on their backs. The frequent falling over onto their backs prevented us from performing classical assays of motor function such as rotor rod or foot printing. Instead, we assayed the righting reflex, open field exploration and USV. We also tested for clinically relevant features such as tremor and dystonia-like postures, which often arise with cerebellar dysfunction. We found that *En1^Cre/+^;Atoh1^fl/-^* mice perform poorly compared to control littermates during the righting reflex. They were significantly slower in returning to the right side when compared to *Atoh1^fl/+^* mice at P8 and P10, and slower than the *En1^Cre/+^;Atoh1^fl/+^*mice only at P10 (**Figure 4B**). Because all mice attempted to turn right-side-up immediately after being placed on their backs, it is likely that this delay in righting is the result of impaired motor coordination rather than an abnormal sense of gravity. Next, we tested whether *En1^Cre/+^;Atoh1^fl/-^* mice showed abnormal USVs when briefly separated from their mothers (**Figure 4C**). We found that call time in *En1^Cre/+^;Atoh1^fl/-^* mice was shorter than those observed in control littermates and that *En1^Cre/+^;Atoh1^fl/-^* mice called less frequently than *Atoh1^fl/+^* mice (**Figure 4C-E**). We next quantified how *En1^Cre/+^;Atoh1^fl/-^* mice moved in an open field (**Figure 4F**). The distance traveled or movement time in a 15-min-period was not significantly impaired (total distance (cm): *Atoh1^fl/+^*: 58.1±11.7; *En1^Cre/+^;Atoh1^fl/+^*: 35.7±9.9; *Atoh1^fl/-^*: 44.4±6.5; *En1^Cre/+^;Atoh1^fl/-^*: 43.1±12.4; Kruskal-Wallis test p=0.32; movement time (s): *Atoh1^fl/+^*: 55.9±5.9; *En1^Cre/+^;Atoh1^fl/+^*: 42.1±7.2; *Atoh1^fl/-^*: 50.1±7.4; *En1^Cre/+^;Atoh1^fl/-^*: 83.5±17.7; Kruskal-Wallis test p=0.14). However, *En1^Cre/+^;Atoh1^fl/-^* mutant mice traveled slower than *Atoh1^fl/+^* control mice and the *En1^Cre/+^;Atoh1^fl/+^* mice made more isolated movements during their trajectory compared to all their littermate controls (**Figure 4F-H**). Finally, we observed a tremor in the mutants and measured the severity with our custom-made tremor monitor (**Figure 4I**) (Brown et al., 2020). We found that *En1^Cre/+^;Atoh1^fl/-^* mice had a higher power tremor in the 12-16 Hz frequency range. This range corresponds to physiological tremor and indicates the presence of a pathophysiological defect that that emerges from a rise in baseline values. The mutant mice also had a higher peak tremor power compared to all control littermates (**Figure 4K**). Together, we uncover that the lack of granule cells in developing *En1^Cre/+^;Atoh1^fl/-^* mice leads to abnormal cerebellar-dependent behaviors.

**Figure 4.**
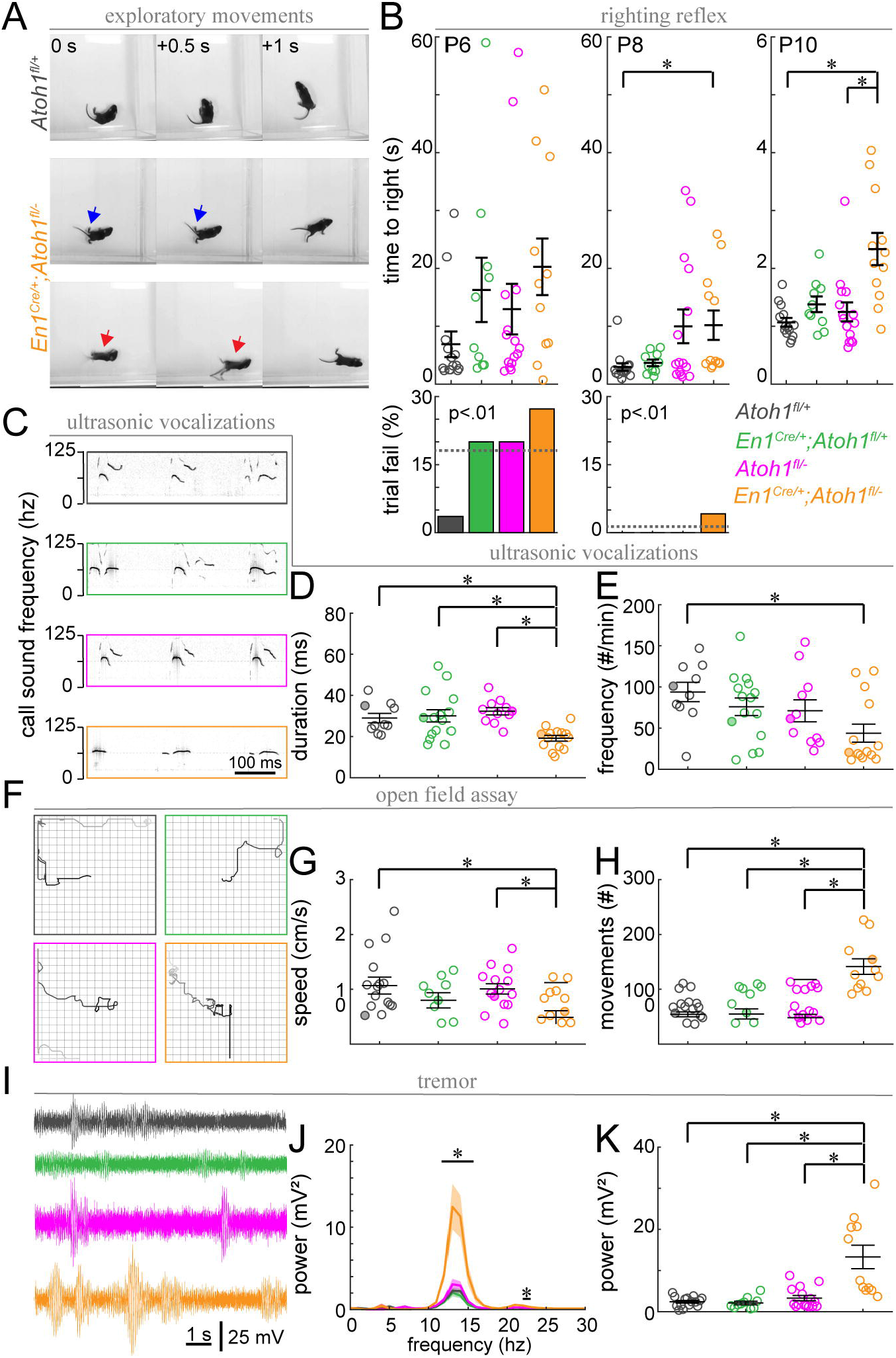
Neonatal *En1^Cre/+^;Atoh1^fl/-^* mice have abnormal motor coordination and vocalization behavior. **A.** Timed-series photos of *Atoh1^fl/+^* (grey) and *En1^Cre/+^;Atoh1^fl/-^* (orange) mice. *En1^Cre/+^;Atoh1^fl/-^* mice have a wide stance (blue arrows) and fall on their backs (red arrows). **B.** Time to right in the righting reflex of P6, P8 and P10 mice (top) and percentage failed trials (bottom). **C.** Representative ultrasonic vocalization traces (filled circles in **D** and **E**). **D.** Duration of vocalizations. **E.** Frequency of vocalizations. **F.** Representative tracks of mice in the open field (filled circles in **G** and **H**). **G.** Movement speed. **H.** Number of movements. **I.** Representative power spectra of tremor. **J.** Tremor power at different frequencies. **K.** Peak tremor power. N-numbers: *Atoh1^fl/+^* (grey): N=10-15; *En1^Cre/+^;Atoh1^fl/+^* (green): N=9-15; *Atoh1^fl/-^* (pink): N=11-15; *En1^Cre/+^;Atoh1^fl/-^* (orange): N=11-14. Significance was determined using a nonparametric Kruskal-Wallis test followed by a Tukey-Kramer post-hoc test. *P<0.05.

## DISCUSSION

In this paper, we used *En1^Cre/+^;Atoh1^fl/-^* mice as a model of cerebellar agranularity to test how cell-to-cell interactions impact the formation of functional circuits. Using this model with circuit-wide loss of granule cell neurogenesis, we uncovered how these late-born cells influence the functional development of their downstream synaptic partners, the Purkinje cells. We find that granule cell elimination halts the anatomical and functional maturation of postnatal Purkinje cells. Notably, granule cell neurogenesis is impaired in premature infants with cerebellar hemorrhages as proliferating granule cell precursors are highly vulnerable to hemorrhages, likely because of their high metabolic demand (Dobbing, 1974; Gano and Barkovich, 2019; Hortensius et al., 2018). The loss of just a few precursors has exponential effects on the number of granule cells that integrate into the cerebellar circuit (Corrales et al., 2004, 2006). Our findings contribute to understanding how early changes in cerebellar volume become correlated with downstream circuit dysfunction and the resulting neurological disorders observed in premature infants (Dijkshoorn et al., 2020).

There are several caveats to using an agranular model to investigate circuitry. For instance, loss of morphogenetic processes that determine cerebellar architecture including its size (Dahmane and Ruiz i Altaba, 1999), foliation (Corrales et al., 2006) and layering (Miyata et al., 2010) complicate interpretations for how Purkinje cells directly respond to granule cells. Our analysis of very young postnatal mice largely addresses this concern. In addition, it is possible that changes in cerebellar shape and size could influence forebrain regions (Kuemerle et al., 2007). Hence, the vast connectivity of the cerebellum with a number of regions such as the hippocampus and prefrontal cortex could contribute to the non-motor vocalization defects we observed (Liu et al., 2020; McAfee et al., 2019). Still, the regional specificity of cerebellar circuitry that mediates non-motor connectivity is likely obscured in *En1^Cre/+^;Atoh1^fl/-^* mice (Badura et al., 2018; Stoodley and Limperopoulos, 2016). Furthermore, the presence of excitatory rhombic lip-derived neurons that escape *Atoh1* deletion, such as the remaining unipolar brush cells (**Supp. 1B-C**), suggests the requirement of manipulating combinatorial molecular domains to fully target the excitatory neuron lineage (Chizhikov et al., 2010; Yeung et al., 2014). Despite these caveats, our mouse model has several advantages over previously described agranular mice when it comes to the contribution of granule cell neurogenesis to cerebellar development and function. Because our model takes advantage of a conditional genetic strategy that only targets the rhombic lip lineage, our manipulation does not affect cell intrinsic developmental Purkinje cell programs (Herrup, 1983; Miyata et al., 2010; Sheldon et al., 1997). Unlike previous models, our approach is independent of procedural variations (Altman and Anderson, 1971; Sathyanesan et al., 2018; Yoo et al., 2014), targets the entire *Atoh1* lineage in the cerebellum (Ben-Arie et al., 1997; Jensen et al., 2002, 2004) and allows us to study postnatal development. As a result, mice with the *En1^Cre/+^;Atoh1^fl/-^* genotype have highly penetrant, consistent and reliable anatomical and functional phenotypes, all of which have provided key insights in how cerebellar lineages shape circuit development and behavior.

The cerebellum controls motor and non-motor behaviors (Hull, 2020; Wagner and Luo, 2020). Regardless of the specific behavior, Purkinje cells are always at the center of the responsible circuit. What is intriguing to us is that during the first two weeks of life in mice, motor capabilities become more precise (Lalonde and Strazielle, 2015), in concert with the refinement of circuitry (White and Sillitoe, 2013). During this period, Purkinje cell innervation switches from climbing fibers and mossy fibers to climbing fibers and parallel fibers (Mason and Gregory, 1984), climbing fibers are pruned and a “winner” establishes a single Purkinje cell target (Kano et al., 2018) and Purkinje cell zones are sharpened (White et al., 2014). Accompanying these changes are emergent properties of the two Purkinje cell spike profiles, simple spikes and complex spikes. In addition to the increased complexity of intrinsic cellular properties (McKay and Turner, 2005), we postulated that intercellular interactions during development may support the maturation of Purkinje cell firing. In control mice, we observed dynamic changes in normal Purkinje cell firing between P7 and P14. We found an increase in firing rate that was not uniformly acquired but was present during bursts of rapid firing that were interspersed with frequent pauses without Purkinje cell action potentials. We previously reported burst-like Purkinje cell firing from P15-19, although by P30 the pattern acquires the regularity that is characteristic of adults (Arancillo et al., 2015). Thus, burst-like firing occurs at intermediate stages of normal Purkinje cell development. Interestingly, bursting Purkinje cell firing patterns are also observed in mouse models of ataxia, tremor, and dystonia (Brown et al., 2020; Fremont et al., 2014; LeDoux and Lorden, 2002; Miterko et al., 2019; White and Sillitoe, 2017; White et al., 2016). The dynamically adapting circuit in controls and the range of disease severity in disease models with bursting Purkinje cells raise the possibility that Purkinje cell firing is differentially decoded by downstream neurons based on the age of the mice. The data also indicate that the intermediate stages of Purkinje cell development not only highlight a developmental phase characterized by erratic neuronal activity, but that this mode of firing represents a pathophysiological hallmark that could be a default network state in different diseases.

The severity of motor impairments and electrophysiological changes in Purkinje cells in our agranular mice was surprising, given that silencing granule cell synapses results in minor changes in Purkinje cell firing (Galliano et al., 2013). Furthermore, in multiple models, impairing parallel fiber synapses results in motor impairments that can be assessed with the rotor rod assay (Aiba et al., 1994; Park et al., 2019), which we could not do due to the severity of motor impairment in *En1^Cre/+^;Atoh1^fl/-^* mice. Taking these results together, from a technical standpoint, when one seeks to resolve developmental mechanisms, we must not only consider *what* is manipulated, but also *how* it is manipulated. As such, the timing of neurogenesis is a primary consideration. Purkinje cells are generated between E10-E13 (Hashimoto and Mikoshiba, 2003) and granule cell progenitors from ~E13 onwards (Machold and Fishell, 2005; Rose et al., 2009; Wang et al., 2005). But, whereas Purkinje cells migrate into the core of the cerebellar anlage upon their birth, granule cells first migrate over the surface of the developing cerebellum and proliferate extensively in the external granular layer to increase the precursor pool (Wingate and Hatten, 1999). Only after this phase do they migrate radially past Purkinje cells, the first potential opportunity for direct cell-to-cell interactions. Based on our data, we argue that the initial communication between Purkinje cells and granule cells sets the efficiency of Purkinje cell function because of a direct influence on the establishment of Purkinje cell spikes. The data further suggest that granule cells shape Purkinje cell development through structural as well as synaptic signals. Thus, insults to granule cell proliferation and an obstruction of granule cell neurogenesis may have different, and perhaps more severe effects on downstream Purkinje cell function, compared to lesions of mature granule cells.

## ACKNOWLEDGEMENTS

This work was supported by Baylor College of Medicine (BCM), Texas Children’s Hospital, The Hamill Foundation, BCM IDDRC U54HD083092 (Neurovisualization Core and the Mouse Neurobehavioral Core), and the National Institutes of Neurological Disorders and Stroke (NINDS) R01NS089664 and R01NS100874 to RVS and F31NS101891 to AMB. HYZ is supported by the Howard Hughes Medical Institute (HHMI).

## AUTHOR CONTRIBUTIONS

MEvdH and RVS conceived the project and wrote the paper; HYZ contributed to discussions that inspired some of the studies. MEvdH, TL, EPL, FI, and RP collected data. MEvdH and AMB analyzed data. All authors interpreted results and edited the final version of the paper.

## METHODS

### Animals

All mice used in this study were housed in a Level 3, AALAS-certified facility. All experiments and studies that involved mice were reviewed and approved by the Institutional Animal Care and Use Committee (IACUC) of Baylor College of Medicine (BCM). The following transgenic mouse lines were used for the preparation of this manuscript: *Atoh1^FlpO^* (van der Heijden and Zoghbi, 2018); *En1^Cre^* (*En1^tm2(cre)Wrst/J^*, JAX:007916); *Ai65* (*Gt(ROSA)26Sor^tm65.1(CAG-tdTomato)Hze^*, JAX:021875); *Atoh1^Flox^* (*Atoh1^tm3Hzo^*, MGI:4420944). Ear tissue or tail clips were collected before weaning and used for genotyping and identification of the different alleles used. For all mice, P0 was defined as the day of birth.

### Tissue processing

Brain and spinal cord tissue was collected as described in our previous publications (Brown et al., 2019; Zhou et al., 2020). First, we anesthetized mice with Avertin. Once the mice did not respond to toe or tail pinch, we accessed the chest cavity and then penetrated the heart with a butterfly needle for perfusions. The mice were perfused with 1M phosphate-buffered saline (PBS pH 7.4) to remove blood from the tissue and 4% paraformaldehyde (PFA) to fix the tissue. The tissue was concomitantly post-fixed overnight in 4% PFA at 4 °C. Tissue was cryoprotected in a sucrose gradient (10% ➕ 20% ➕ 30% sucrose in PBS) at 4 °C, each step lasting until the tissue sank to the bottom of a 15 mL tube. Tissue was frozen in optimal cutting temperature (OCT) solution and stored at −80 °C until cut. All tissue was cut into 40 μm free-floating sections and stored in PBS at 4 °C until it was used for immunohi stochemi stry.

### Immunohistochemistry

Free floating sections were stained according to the following protocol. Free floating sections were blocked in 10% normal goat or donkey serum and 0.1% Triton-X in PBS for two hours. Next, sections were incubated overnight in primary antibody in blocking solution. Tissue was washed three times for five minutes in 0.1% Triton-X in PBS (PBS-T). For fluorescent staining, the tissue was incubated for two hours in PBS-T with preferred secondary antibody conjugated to an Alexa fluorophore. Finally, sections were washed three times in PBS-T and mounted on electrostatically coated slides with hard-set, DAPI containing mounting medium. Alternatively, for DAB staining, we incubated the tissue for two hours in PBS-T with the preferred secondary antibody that was conjugated to HRP. After washing three times in PBS-T, the tissue was incubated with DAB solution until the desired color intensity was reached. The DAB color reaction was stopped by washing tissue three times with PBS-T. The tissue was then mounted on electrostatically coated glass slides, dehydrated in an ethanol series (70% ➕ 90% ➕ 100%) and then mounted using Xylene or histoclear. All steps of immunohistochemistry were performed at room temperature. All mounted slides were stored at 4 °C until they were imaged.

The following primary antibodies were used for the data described in this manuscript: guinea pig (gp)-α-Calbindin (1:1,000; SySy; #214004); rabbit (rb)-α-gamma-aminobutyric acid receptor α6 (GABARα6; 1:500; Millipore Sigma; #AB5610), rb-α-T-box brain protein 2 (Tbr2; 1:500; Abcam; #AB23345), mouse (ms)-α-Calretinin (1:500; Swant; #6B3); ms-α-Neurofilament Heavy (NFH; 1:1,000; Biolegend; #801701); rb-α-Hyperpolarization Activated Cyclic Nucleotide Gated Potassium Channel 1 (HCN1; 1:500; Alomone Lab; #APC-056); goat (gt)-α-RAR-related orphan receptor alpha (RORα; 1:250; Santa Cruz; #F2510); rb-α-parvalbumin (PV; 1:1,000; Swant; #PV25); rb-α-neurogranin (1:500; Millipore Sigma; #AB5620); ms-α-ZebrinII (1:500; kind gift from Dr. Richard Hawkes, University of Calgary, Calgary, Alberta, Canada); rb-α-Vglut1 (1:500; SySy; #135302); rb-α-Vglut2 (1:500; SySy; #135403). The following secondary antibodies were used for immunohistochemistry: HRP-conjugated goat (gt)-α-mouse; gt-α-rabbit; and rabbit (dk)-α-goat (1:200; DAKO). The following secondary antibodies were used for immunofluorescence: dk-α-mouse IgG Alexa Fluor 488 (1; 1,500; Thermo Fisher Scientific; #A21202) and gt-α-gp IgG Alexa Fluor 488 (1; 1,500; Thermo Fisher Scientific;#A11073).

### Cresyl violet staining

Brain sections were mounted on electrostatically coated glass slides and then dried overnight. Slides were submerged in 100% histoclear and rehydrated in an ethanol series (100% ➕ 90% ➕ 70%). Then, the slides were submerged in cresyl violet solution for staining until sufficiently dark and then dehydrated in an ethanol series (70% ➕ 90% ➕ 100%). Finally, the slides were sealed with a coverslip using Cytoseal mounting media. All steps were performed at room temperature and the mounted slides were stored at 4 °C until they were imaged.

### Neuroanatomical anterograde tracing

Anterograde neuroanatomical tracing of mossy fibers to the cerebellum were performed as described previously (Lackey and Sillitoe, 2020; Sillitoe, 2016). P12 pups were anesthetized with isoflurane on a surgery rig. Hair was removed and an incision was made in the skin over the lower thoracic/upper lumbar spinal cord, using the curvature of the spine as a guide. We used a Nanoject II to inject 0.2 – 1 μl of 2% WGA-Alexa Fluor 555 (Thermo Fisher Scientific; #W32464) and 0.5% Fast Green (Sigma-Aldrich; #F7252, used for visualization) diluted in 0.1 M phosphate-buffered saline (PBS; Sigma-Aldric; #P4417; pH 7.4). Tracers were injected 1 mm below the surface of the spinal cord, on the right side of the dorsal spinal vein. After tracer injection, we applied antibiotic ointment and closed the incision using VetBond (3M; #1469SB) and wound clips (Fine Science Tools; #12032-07). Pups were placed back with the mom after waking up from anesthesia. We placed soft food and hydrogel on the floor cage and monitored closely whether mom was taking care and feeding the pups. Tissue was collected (see section on *Tissue processing* above) for tracer visualization 2 days after the surgery, at P14.

### Golgi-Cox staining

Golgi-Cox staining was performed according to previously described protocols (Brown et al., 2019) and manufacturer’s instructions (FD Neurotechnologies; #PK401). Brains were dissected from the skulls and immediately emerged in the staining solution. After staining, sections were cut in 10 μm thickness and directly mounted on electrostatically coated slides. Tissue was then dehydrated in an ethanol series (70% ➕ 90% ➕ 100%), cleared with Xylene, and mounted with cytoseal. All slides were dried overnight before imaging and were kept at 4 °C for storage.

### Fluorescence imaging and analysis of staining

Photomicrographs of stained whole mount cerebella and DAB stained cerebellar sections were acquired using Leica cameras DPC365FX and DMC2900, respectively, attached to a Leica DM4000 B LED microscope. Photomicrographs of Golgi-Cox stained sections and WGA-555 tracing were made using Zeiss cameras AxioCam MRc5 and AxiaCam Mrm, respectively, attached to a Zeiss Axio Imager.M2 microscope. Whole mount images were stitched together using Adobe Photoshop. Color brightness and contrast were adjusted using ImageJ. Photomicrographs of images were cropped to desired size using Adobe Illustrator. Sholl analysis was performed using the build-in Sholl analysis module in ImageJ (Ferreira et al., 2014) and false-positive intersections were manually subtracted from the counts.

### In vivo electrophysiology

All *in vivo*, anesthetized experiments were performed as described in previous publications (Arancillo et al., 2015; White and Sillitoe, 2017). Specifically, we anesthetized mice using a mixture of ketamine 80 (mg/kg) and dexmedetomidine (16 mg/kg). We held mice on a heated surgery pad. We removed hair from skull and made an incision in the skin over the anterior part of the skull. We stabilized the heads of our mice using ear bars and a mouth mount when animals were large enough (most P11-P14 mice) and otherwise glued the mouse skull (P7-P10 mice) to a plastic mount that was attached to ear bars on our stereotaxic surgery rig to stabilize the head during recordings. Using a sharp needle or dental drill, we made a craniotomy in the interparietal bone plate, ~3 mm dorsal from lambda and ~3 mm lateral from the midline, with a diameter of ~3mm. We kept our surgical coordinates consistent based on the distance from lambda across mice of all ages, as the skull undergoes significant growth during the ages at which we measured neural activity. After making a craniotomy, we recorded neural activity using tungsten electrodes (Thomas Recording, Germany) and then the digitized the signals into Spike2 (CED, England). We recorded neural activity from cells that were 0-2 mm below the brain surface.

### Analysis of in vivo electrophysiological recordings

All electrophysiological recording data were spike sorted in Spike2. We sorted out three types of spikes: simple spikes, complex spikes, and doublets. Complex spikes were characterized by their large amplitude, and post-spike depolarization and smaller wavelets. Doublets were characterized as action potentials that were followed by one or more smaller action potentials within 20 ms after the initial action potential. All other action potentials were characterized as simple spikes (see examples in **Figure 3** and **Supp. F3**). We only included traces with clearly identifiable complex spikes or doublets and analyzed only cells from which we could obtain a sufficient long and stable recording (186 ± 6.5 s; minimum = 75s) with an optimal signal to noise rAll mice used in this study were hoatio.

After spike sorting our traces in Spike 2, we analyzed the frequency and regularity of firing patterns in MATLAB. For this study, we defined frequency as number of all spikes observed in the total analyzed recording time (spikes/s). Our measures of global regularity or burstiness (CV) and regularity (CV2) were based on the interspike intervals (ISI) between two adjacent spikes (in s). CV = stdev(ISI)/mean(ISI), and CV2 = mean(2*|ISIn-ISIn-1|/ (ISIn+ISIn-1)). Pause Percent was the proportion of the recording time during which the ISI was longer than five times the mean ISI for each independent cell, defined as followed: (sum(ISI>5*mean(ISI)))/(total recording time).

### Behavioral analyses

Righting reflex was measured on P6, P8, and P10 as followed. Mouse was placed on its back in a clean cage without bedding. One finger was used to stabilize the mouse on its back. The timer was set the moment the experimenter removed their finger, and time was recorded until mouse righted itself up to four paws. All mice were tested twice on each time point. A “failed” trial was defined when the mouse did not right itself within one minute (sixty seconds).

At P7, we recorded pup vocalizations as described previously (Yin et al., 2018). Pups were placed in an anechoic, sound-attenuating chamber (Med Associates Inc.). The pup was placed in a round plastic tub that was positioned near a CM16 microphone (Avisoft Bioacoustics) that was located in the center of the chamber. Sound was amplified and digitized using UltraSoundGate 416H at a 250 kHz sampling rate and bit depth of 16. Avisoft RECORDER software was used to collect the recordings. Ultrasonic vocalizations were monitored for 2 min for each animal.

We also performed an open field assay at P13 as previously described (Alcott et al., 2020). Mice were habituated to a room with light set to 200 lux and ambient white noise to 60 dB. We placed each mouse in the center of an open field (40×40×30 cm chamber). The chamber has photobeams that could record movement. Each mouse was tested for 15 minutes and activity was recorded using Fusion software (Accuscan Instruments). We analyzed the data for total distance traveled, movement time, speed, and total movements during the 15-minute test period.

We measured tremor using our custom-built tremor monitor (Brown et al., 2020). Each mouse was placed in the tremor chamber, which is a translucent box with an open top that is suspended in the air by eight elastic cords that are attached to four metal rods. An accelerometer is attached to the bottom of the box. Mice were allowed to habituate to the chamber for 120 second prior to tremor recordings. The mice are free to move around in the box. Power spectrums of tremor were assessed using Fast Fourier transform (FFT) with Hanning window in Spike2 software as previously described (Brown et al., 2020). FFT frequency was target to ~1 Hz per bin.

### Statistical analysis

Statistical analyses were performed in MATLAB. For electrophysiology data, analysis was performed using a t-test. We used a Bonferroni-corrected p-value for statistical significance to avoid false positives (p<0.000625 (=0.05/8) was accepted as statistically significant). We performed the hierarchical cluster analysis on the first three principle components of the firing frequency, frequency mode, CV, CV2, and Pause Percent. For behavioral data, we performed a Kruskal-Wallis test followed by a Tukey-Kramer post-hoc test to define significance between independent groups. For these tests, we accepted p<0.05 as statistically significant.

**Supplemental Figure 1 – with Figure 1. Conditional deletion of *Atoh1* from the *En1* domain targets excitatory, but not inhibitory cerebellar cell types. A.** *En1^Cre/+^;Atoh1^fl/-^* mice lack differentiated granule cells, identified with GABARα6. **B.** and **C.** *En1^Cre/+^;Atoh1^fl/-^* mice have a reduction in unipolar brush cells, identified by Calretinin and Tbr2, respectively. **D.** *En1^Cre/+^;Atoh1^fl/-^* mice have dense staining for NFH-positive cells that mark Purkinje cells and excitatory nuclei (interposed nucleus shown here). **E** through **H.** *En1^Cre/+^;Atoh1^fl/-^* mice have a high density fot inhibitory neurons, revealed with the expression of RORα (**E**), HCN1 (**F**), Neurogranin (**G**), and PV (**H**). All images are representative for N=3 brains for each genotype.

**Supplemental Figure 2 – with Figure 2. *En1^Cre/+^;Atoh1^fl/-^* mice express mature cerebellum stripe markers, but do not form clear-cut stripes. A.** Top view of a control P14 brain. Dotted line shows the position of schematic in **B.** and where the section in **C**. was taken from. **B.** Schematic of Purkinje cell ZebrinII (pink) and PLCβ4 (green) patterns in a control section illustrating the striped patterns at P14. **C.** Staining of ZebrinII (pink) and PLCβ4 (green). **D.-G.** Higher power magnification images of insets in **C. H.** Top view of *En1^Cre/+^;Atoh1^fl/-^* P14 brain. Dotted line shows position of schematic in **I.** and where the section in **J**. is taken from. **I.** Schematic of Purkinje cell ZebrinII (pink) and PLCβ4 (green) patterns in *En1^Cre/+^;Atoh1^fl/-^* mice showing a clustered pattern at P14. **J.** Staining of ZebrinII (pink) and PLCβ4 (green). **K-L.** Higher power magnification images of insets in **J.** All images are representative of N=3 brains per genotype.

**Supplemental Figure 3 – with Figure 3. Representative traces of in vivo firing properties of Purkinje cells from P7-P14 control and P14 *En1^Cre/+^;Atoh1^fl/-^* mice.** Left, representative trace is 10 s long. Middle, trace is 1 second long. Right, representative simple spike and doublet or complex spike (depending on most prevalent spike type). Representative cells were chosen based on that particular cell’s firing properties being closest to the group average.

**Video 1 – with Figure 4. *En1^Cre/+^;Atoh1^fl/-^* mice have visible motor impairments.** At P14, control mice explore the open box smoothly, whereas *En1^Cre/+^;Atoh1^fl/-^* mice have a visible tremor, often fall on their backs, and have dystonia-like muscle contractions in their hind paws.

## REFERENCES

Aiba, A., Kano, M., Chen, C., Stanton, M.E., Fox, G.D., Herrup, K., Zwingman, T.A., and Tonegawa, S. (1994). Deficient cerebellar long-term depression and impaired motor learning in mGluR1 mutant mice. Cell 79, 377–388.

Alcott, C.E., Yalamanchili, H.K., Ji, P., van der Heijden, M.E., Saltzman, A., Elrod, N., Lin, A., Leng, M., Bhatt, B., Hao, S., et al. (2020). Partial loss of CFIm25 causes learning deficits and aberrant neuronal alternative polyadenylation. Elife 9.

Altman, J., and Anderson, W.J. (1971). Irradiation of the cerebellum in infant rats with low-level x-ray: histological and cytological effects during infancy and adulthood. Exp. Neurol. 30, 492–509.

Arancillo, M., White, J.J., Lin, T., Stay, T.L., and Sillitoe, R.V. (2015). In vivo analysis of Purkinje cell firing properties during postnatal mouse development. J. Neurophysiol. 113, 578–591.

Badura, A., Verpeut, J.L., Metzger, J.W., Pereira, T.D., Pisano, T.J., Deverett, B., Bakshinskaya, D.E., and Wang, S.S.-H. (2018). Normal cognitive and social development require posterior cerebellar activity. Elife 7.

Ben-Arie, N., Bellen, H.J., Armstrong, D.L., McCall, A.E., Gordadze, P.R., Guo, Q., Matzuk, M.M., and Zoghbi, H.Y. (1997). Math1 is essential for genesis of cerebellar granule neurons. Nature 390, 169–172.

Bradley, P., and Berry, M. (1976). The effects of reduced climbing and parallel fibre input on Purkinje cell dendritic growth. Brain Res. 109, 133–151.

Brochu, G., Maler, L., and Hawkes, R. (1990). Zebrin II: a polypeptide antigen expressed selectively by Purkinje cells reveals compartments in rat and fish cerebellum. J. Comp. Neurol. 291, 538–552.

Brown, A.M., Arancillo, M., Lin, T., Catt, D.R., Zhou, J., Lackey, E.P., Stay, T.L., Zuo, Z., White, J.J., and Sillitoe, R.V. (2019). Molecular layer interneurons shape the spike activity of cerebellar Purkinje cells. Sci. Rep. 9, 1742.

Brown, A.M., White, J.J., van der Heijden, M.E., Zhou, J., Lin, T., and Sillitoe, R.V. (2020). Purkinje cell misfiring generates high-amplitude action tremors that are corrected by cerebellar deep brain stimulation. Elife 9.

Chang, C.H., Chang, F.M., Yu, C.H., Ko, H.C., and Chen, H.Y. (2000). Assessment of fetal cerebellar volume using three-dimensional ultrasound. Ultrasound Med Biol 26, 981–988.

Chizhikov, V.V., Lindgren, A.G., Mishima, Y., Roberts, R.W., Aldinger, K.A., Miesegaes, G.R., Currle, D.S., Monuki, E.S., and Millen, K.J. (2010). Lmx1a regulates fates and location of cells originating from the cerebellar rhombic lip and telencephalic cortical hem. Proc. Natl. Acad. Sci. USA 107, 10725–10730.

Corrales, J.D., Rocco, G.L., Blaess, S., Guo, Q., and Joyner, A.L. (2004). Spatial pattern of sonic hedgehog signaling through Gli genes during cerebellum development. Development 131, 5581–5590.

Corrales, J.D., Blaess, S., Mahoney, E.M., and Joyner, A.L. (2006). The level of sonic hedgehog signaling regulates the complexity of cerebellar foliation. Development 133, 1811–1821.

Dahmane, N., and Ruizi Altaba, A. (1999). Sonic hedgehog regulates the growth and patterning of the cerebellum. Development 126, 3089–3100.

Davie, J.T., Clark, B.A., and Häusser, M. (2008). The origin of the complex spike in cerebellar Purkinje cells. J. Neurosci. 28, 7599–7609.

Davis, C.A., and Joyner, A.L. (1988). Expression patterns of the homeo box-containing genes En-1 and En-2 and the proto-oncogene int-1 diverge during mouse development. Genes Dev. 2, 1736–1744.

Dijkshoorn, A.B.C., Turk, E., Hortensius, L.M., van der Aa, N.E., Hoebeek, F.E., Groenendaal, F., Benders, M.J.N.L., and Dudink, J. (2020). Preterm infants with isolated cerebellar hemorrhage show bilateral cortical alterations at term equivalent age. Sci. Rep. 10, 5283.

Dobbing, J. (1974). The later growth of the brain and its vulnerability. Pediatrics 53, 2–6.

Dusart, I., Guenet, J.L., and Sotelo, C. (2006). Purkinje cell death: differences between developmental cell death and neurodegenerative death in mutant mice. Cerebellum 5, 163–173.

Fremont, R., Calderon, D.P., Maleki, S., and Khodakhah, K. (2014). Abnormal high-frequency burst firing of cerebellar neurons in rapid-onset dystonia-parkinsonism. J. Neurosci. 34, 11723–11732.

Fujita, E., Tanabe, Y., Shiota, A., Ueda, M., Suwa, K., Momoi, M.Y., and Momoi, T. (2008). Ultrasonic vocalization impairment of Foxp2 (R552H) knockin mice related to speech-language disorder and abnormality of Purkinje cells. Proc. Natl. Acad. Sci. USA 105, 3117–3122.

Fujita, H., Morita, N., Furuichi, T., and Sugihara, I. (2012). Clustered fine compartmentalization of the mouse embryonic cerebellar cortex and its rearrangement into the postnatal striped configuration. J. Neurosci. 32, 15688–15703.

Galliano, E., Gao, Z., Schonewille, M., Todorov, B., Simons, E., Pop, A.S., D’Angelo, E., van den Maagdenberg, A.M.J.M., Hoebeek, F.E., and De Zeeuw, C.I. (2013). Silencing the majority of cerebellar granule cells uncovers their essential role in motor learning and consolidation. Cell Rep. 3, 1239–1251.

Gano, D., and Barkovich, A.J. (2019). Cerebellar hypoplasia of prematurity: Causes and consequences. Handb Clin Neurol 162, 201–216.

Gebre, S.A., Reeber, S.L., and Sillitoe, R.V. (2012). Parasagittal compartmentation of cerebellar mossy fibers as revealed by the patterned expression of vesicular glutamate transporters VGLUT1 and VGLUT2. Brain Struct. Funct. 217, 165–180.

Gold, D.A., Gent, P.M., and Hamilton, B.A. (2007). ROR alpha in genetic control of cerebellum development: 50 staggering years. Brain Res. 1140, 19–25.

Hashimoto, M., and Mikoshiba, K. (2003). Mediolateral compartmentalization of the cerebellum is determined on the “birth date” of Purkinje cells. J. Neurosci. 23, 11342–11351.

Herrup, K. (1983). Role of staggerer gene in determining cell number in cerebellar cortex. I. Granule cell death is an indirect consequence of staggerer gene action. Developmental Brain Research 11, 267–274.

Hortensius, L.M., Dijkshoorn, A.B.C., Ecury-Goossen, G.M., Steggerda, S.J., Hoebeek, F.E., Benders, M.J.N.L., and Dudink, J. (2018). Neurodevelopmental consequences of preterm isolated cerebellar hemorrhage: A systematic review. Pediatrics 142.

Hoshino, M., Nakamura, S., Mori, K., Kawauchi, T., Terao, M., Nishimura, Y.V., Fukuda, A., Fuse, T., Matsuo, N., Sone, M., et al. (2005). Ptf1a, a bHLH transcriptional gene, defines GABAergic neuronal fates in cerebellum. Neuron 47, 201–213.

Huang, C., Gammon, S.J., Dieterle, M., Huang, R.H., Likins, L., and Ricklefs, R.E. (2014). Dramatic increases in number of cerebellar granule-cell-Purkinje-cell synapses across several mammals. Mammalian Biology 79, 163–169.

Hull, C. (2020). Prediction signals in the cerebellum: beyond supervised motor learning. Elife 9.

Iskusnykh, I.Y., Buddington, R.K., and Chizhikov, V.V. (2018). Preterm birth disrupts cerebellar development by affecting granule cell proliferation program and Bergmann glia. Exp. Neurol. 306, 209–221.

Jensen, P., Zoghbi, H.Y., and Goldowitz, D. (2002). Dissection of the cellular and molecular events that position cerebellar Purkinje cells: a study of the math1 null-mutant mouse. J. Neurosci. 22, 8110–8116.

Jensen, P., Smeyne, R., and Goldowitz, D. (2004). Analysis of cerebellar development in math1 null embryos and chimeras. J. Neurosci. 24, 2202–2211.

Kano, M., Watanabe, T., Uesaka, N., and Watanabe, M. (2018). Multiple phases of climbing fiber synapse elimination in the developing cerebellum. Cerebellum 17, 722–734.

Kuemerle, B., Gulden, F., Cherosky, N., Williams, E., and Herrup, K. (2007). The mouse Engrailed genes: a window into autism. Behav. Brain Res. 176, 121–132.

Lackey, E.P., and Sillitoe, R.V. (2020). Eph/ephrin function contributes to the patterning of spinocerebellar mossy fibers into parasagittal zones. Front. Syst. Neurosci.

Lackey, E.P., Heck, D.H., and Sillitoe, R.V. (2018). Recent advances in understanding the mechanisms of cerebellar granule cell development and function and their contribution to behavior. [version 1; peer review: 3 approved]. F1000Res. 7.

Lalonde, R., and Strazielle, C. (2015). Behavioral effects of neonatal lesions on the cerebellar system. Int. J. Dev. Neurosci. 43, 58–65.

Larsell, O. (1952). The morphogenesis and adult pattern of the lobules and fissures of the cerebellum of the white rat. J. Comp. Neurol. 97, 281–356.

LeDoux, M.S., and Lorden, J.F. (2002). Abnormal spontaneous and harmaline-stimulated Purkinje cell activity in the awake genetically dystonic rat. Exp. Brain Res. 145, 457–467.

Limperopoulos, C., Soul, J.S., Gauvreau, K., Huppi, P.S., Warfield, S.K., Bassan, H., Robertson, R.L., Volpe, J.J., and du Plessis, A.J. (2005). Late gestation cerebellar growth is rapid and impeded by premature birth. Pediatrics 115, 688–695.

Limperopoulos, C., Bassan, H., Gauvreau, K., Robertson, R.L., Sullivan, N.R., Benson, C.B., Avery, L., Stewart, J., Soul, J.S., Ringer, S.A., et al. (2007). Does cerebellar injury in premature infants contribute to the high prevalence of long-term cognitive, learning, and behavioral disability in survivors? Pediatrics 120, 584–593.

Liu, Y., McAfee, S.S., Sillitoe, R.V., and Heck, D.H. (2020). Cerebellar modulation of gamma coherence between prefrontal cortex and hippocampus during spatial working memory decision making. BioRxiv.

Machold, R., and Fishell, G. (2005). Math1 is expressed in temporally discrete pools of cerebellar rhombic-lip neural progenitors. Neuron 48, 17–24.

Mason, C.A., and Gregory, E. (1984). Postnatal maturation of cerebellar mossy and climbing fibers: transient expression of dual features on single axons. J. Neurosci. 4, 1715–1735.

McAfee, S.S., Liu, Y., Sillitoe, R.V., and Heck, D.H. (2019). Cerebellar lobulus simplex and crus I differentially represent phase and phase difference of prefrontal cortical and hippocampal oscillations. Cell Rep. 27, 2328–2334.e3.

McKay, B.E., and Turner, R.W. (2005). Physiological and morphological development of the rat cerebellar Purkinje cell. J. Physiol. (Lond.) 567, 829–850.

Miterko, L.N., White, J.J., Lin, T., Brown, A.M., O’Donovan, K.J., and Sillitoe, R.V. (2019). Persistent motor dysfunction despite homeostatic rescue of cerebellar morphogenesis in the Car8 waddles mutant mouse. Neural Dev. 14, 6.

Miyata, T., Ono, Y., Okamoto, M., Masaoka, M., Sakakibara, A., Kawaguchi, A., Hashimoto, M., and Ogawa, M. (2010). Migration, early axonogenesis, and Reelin-dependent layer-forming behavior of early/posterior-born Purkinje cells in the developing mouse lateral cerebellum. Neural Dev. 5, 23.

Park, H., Kim, T., Kim, J., Yamamoto, Y., and Tanaka-Yamamoto, K. (2019). Inputs from Sequentially Developed Parallel Fibers Are Required for Cerebellar Organization. Cell Rep. 28, 2939–2954.e5.

Puro, D.G., and Woodward, D.J. (1977). Maturation of evoked climbing fiber input to rat cerebellar purkinje cells (I.). Exp. Brain Res. 28, 85–100.

Raman, I.M., and Bean, B.P. (1999). Ionic currents underlying spontaneous action potentials in isolated cerebellar Purkinje neurons. J. Neurosci. 19, 1663–1674.

Rose, M.F., Ahmad, K.A., Thaller, C., and Zoghbi, H.Y. (2009). Excitatory neurons of the proprioceptive, interoceptive, and arousal hindbrain networks share a developmental requirement for Math1. Proc. Natl. Acad. Sci. USA 106, 22462–22467.

Sathyanesan, A., Kundu, S., Abbah, J., and Gallo, V. (2018). Neonatal brain injury causes cerebellar learning deficits and Purkinje cell dysfunction. Nat. Commun. 9, 3235.

Sathyanesan, A., Zhou, J., Scafidi, J., Heck, D.H., Sillitoe, R.V., and Gallo, V. (2019). Emerging connections between cerebellar development, behaviour and complex brain disorders. Nat. Rev. Neurosci. 20, 298–313.

Sheldon, M., Rice, D.S., D’Arcangelo, G., Yoneshima, H., Nakajima, K., Mikoshiba, K., Howell, B.W., Cooper, J.A., Goldowitz, D., and Curran, T. (1997). Scrambler and yotari disrupt the disabled gene and produce a reeler-like phenotype in mice. Nature 389, 730–733.

Sillitoe, R.V. (2016). Mossy fibers terminate directly within purkinje cell zones during mouse development. Cerebellum 15, 14–17.

Sillitoe, R.V., and Hawkes, R. (2002). Whole-mount immunohistochemistry: a high-throughput screen for patterning defects in the mouse cerebellum. J. Histochem. Cytochem. 50, 235–244.

Sokoloff, G., Plumeau, A.M., Mukherjee, D., and Blumberg, M.S. (2015). Twitch-related and rhythmic activation of the developing cerebellar cortex. J. Neurophysiol. 114, 1746–1756.

Steggerda, S.J., Leijser, L.M., Wiggers-de Bruïne, F.T., van der Grond, J., Walther, F.J., and van Wezel-Meijler, G. (2009). Cerebellar injury in preterm infants: incidence and findings on US and MR images. Radiology 252, 190–199.

Stoodley, C.J., and Limperopoulos, C. (2016). Structure-function relationships in the developing cerebellum: Evidence from early-life cerebellar injury and neurodevelopmental disorders. Semin Fetal Neonatal Med 21, 356–364.

Sugihara, I., and Fujita, H. (2013). Peri-and postnatal development of cerebellar compartments in the mouse. Cerebellum 12, 325–327.

van der Heijden, M.E., and Zoghbi, H.Y. (2018). Loss of Atoh1 from neurons regulating hypoxic and hypercapnic chemoresponses causes neonatal respiratory failure in mice. Elife 7.

Volpe, J.J. (2009). Cerebellum of the premature infant: rapidly developing, vulnerable, clinically important. J. Child Neurol. 24, 1085–1104.

Wagner, M.J., and Luo, L. (2020). Neocortex-Cerebellum Circuits for Cognitive Processing. Trends Neurosci. 43, 42–54.

Wang, V.Y., Rose, M.F., and Zoghbi, H.Y. (2005). Math1 expression redefines the rhombic lip derivatives and reveals novel lineages within the brainstem and cerebellum. Neuron 48, 31–43.

White, J.J., and Sillitoe, R.V. (2013). Development of the cerebellum: from gene expression patterns to circuit maps. Wiley Interdiscip Rev Dev Biol 2, 149–164.

White, J.J., and Sillitoe, R.V. (2017). Genetic silencing of olivocerebellar synapses causes dystonia-like behaviour in mice. Nat. Commun. 8, 14912.

White, J.J., Arancillo, M., Stay, T.L., George-Jones, N.A., Levy, S.L., Heck, D.H., and Sillitoe, R.V. (2014). Cerebellar zonal patterning relies on Purkinje cell neurotransmission. J. Neurosci. 34, 8231–8245.

White, J.J., Arancillo, M., King, A., Lin, T., Miterko, L.N., Gebre, S.A., and Sillitoe, R.V. (2016). Pathogenesis of severe ataxia and tremor without the typical signs of neurodegeneration. Neurobiol. Dis. 86, 86–98.

Wingate, R.J., and Hatten, M.E. (1999). The role of the rhombic lip in avian cerebellum development. Development 126, 4395–4404.

Wurst, W., Auerbach, A.B., and Joyner, A.L. (1994). Multiple developmental defects in Engrailed-1 mutant mice: an early mid-hindbrain deletion and patterning defects in forelimbs and sternum. Development 120, 2065–2075.

Yeung, J., Ha, T.J., Swanson, D.J., Choi, K., Tong, Y., and Goldowitz, D. (2014). Wls provides a new compartmental view of the rhombic lip in mouse cerebellar development. J. Neurosci. 34, 12527–12537.

Yin, J., Chen, W., Chao, E.S., Soriano, S., Wang, L., Wang, W., Cummock, S.E., Tao, H., Pang, K., Liu, Z., et al. (2018). Otud7a knockout mice recapitulate many neurological features of 15q13.3 microdeletion syndrome. Am. J. Hum. Genet. 102, 296–308.

Yoo, J.Y.J., Mak, G.K., and Goldowitz, D. (2014). The effect of hemorrhage on the development of the postnatal mouse cerebellum. Exp. Neurol. 252, 85–94.

Zagha, E., Lang, E.J., and Rudy, B. (2008). Kv3.3 channels at the Purkinje cell soma are necessary for generation of the classical complex spike waveform. J. Neurosci. 28, 1291–1300.

Zayek, M.M., Benjamin, J.T., Maertens, P., Trimm, R.F., Lal, C.V., and Eyal, F.G. (2012). Cerebellar hemorrhage: a major morbidity in extremely preterm infants. J. Perinatol. 32, 699–704.

Zhou, Y. (Joy), Brown, A.M., Lackey, E.P., Arancillo, M., Lin, T., and Sillitoe, R.V. (2020). Purkinje cell neurotransmission patterns cerebellar basket cells into zonal modules that are defined by distinct pinceau sizes. BioRxiv.

