## Supplementary figures and images for "Maturation of Purkinje cell firing properties relies on granule cell neurogenesis"

### Supplemental Figure 1

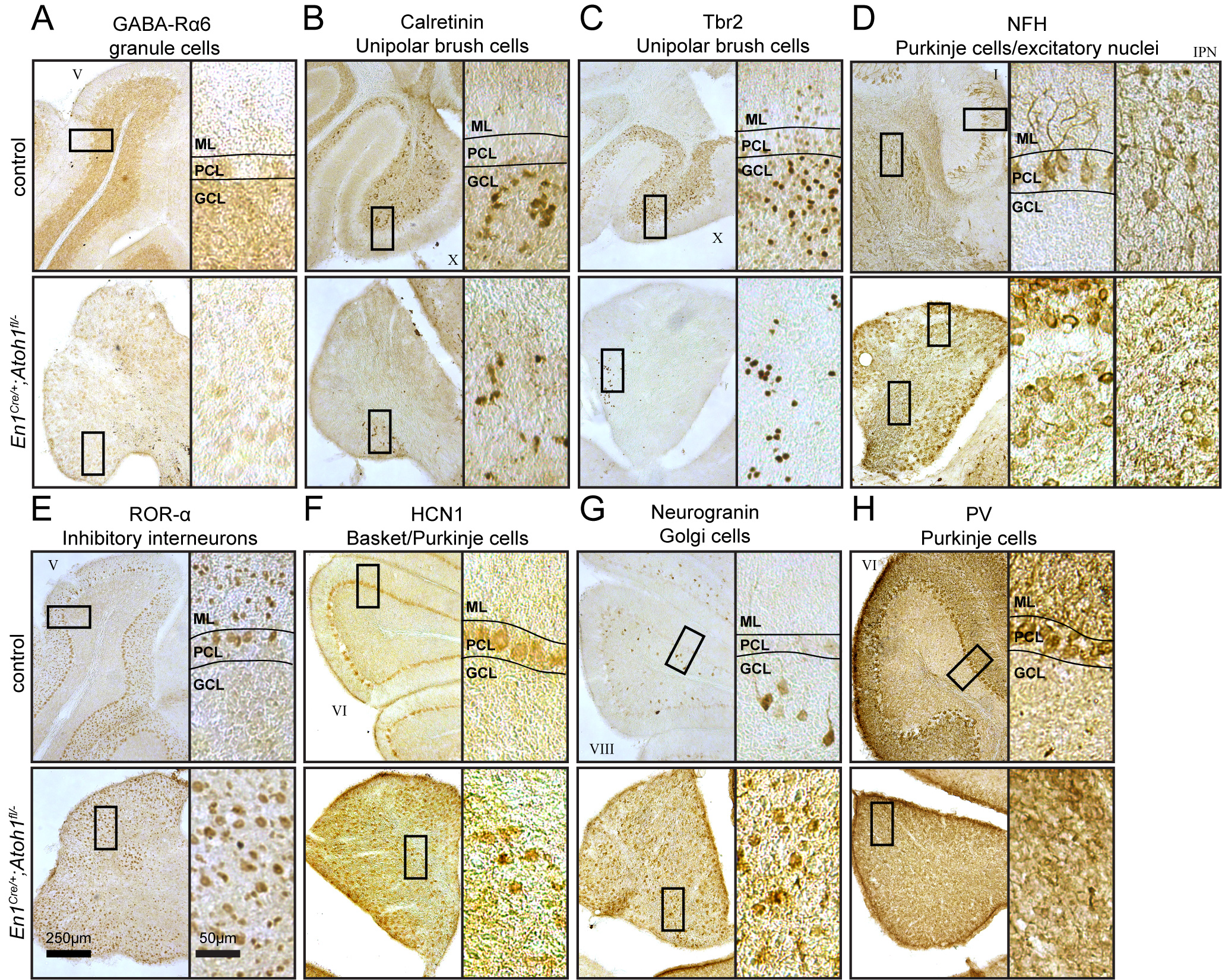

### Supplemental Figure 2

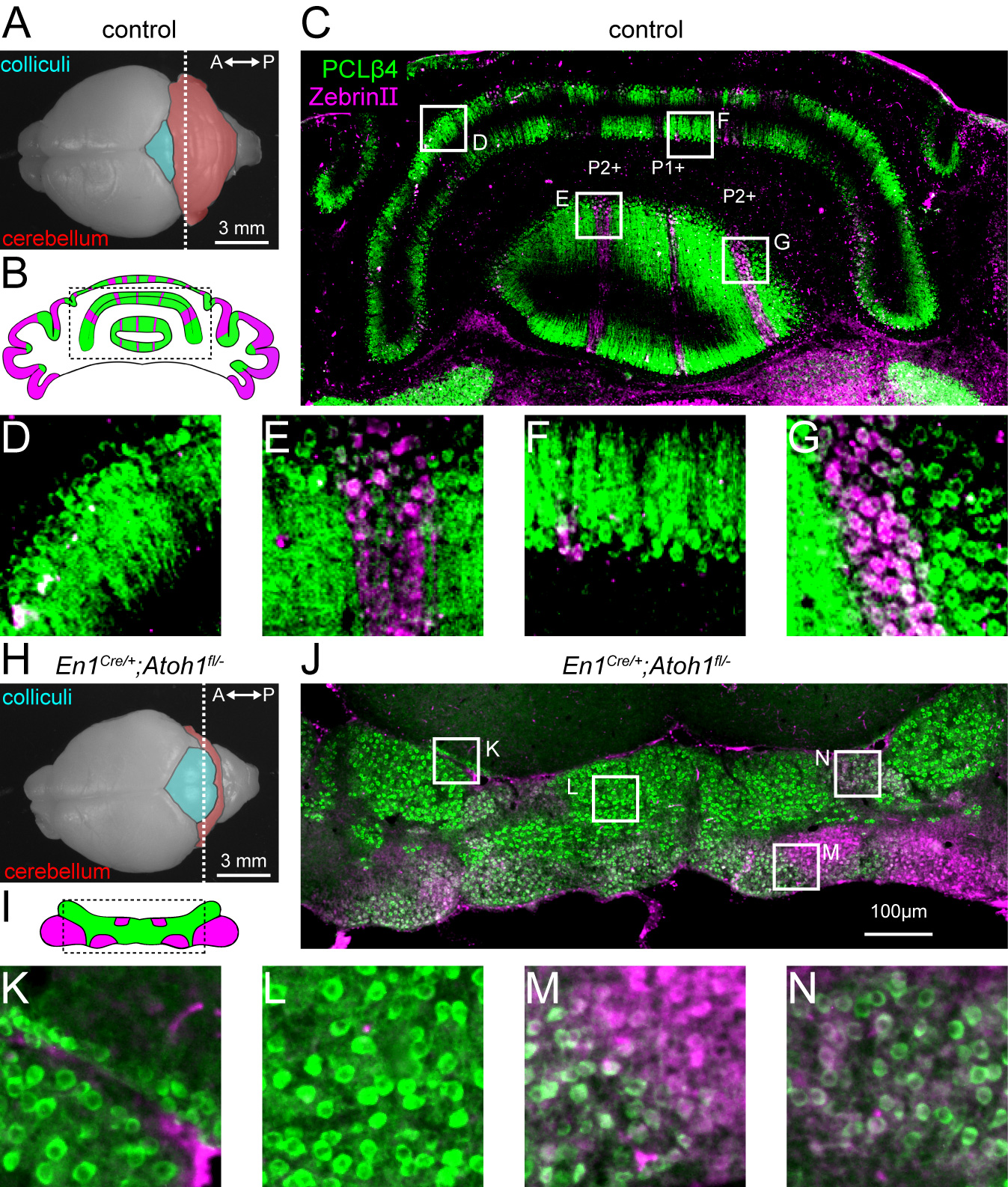

### Supplemental Figure 3

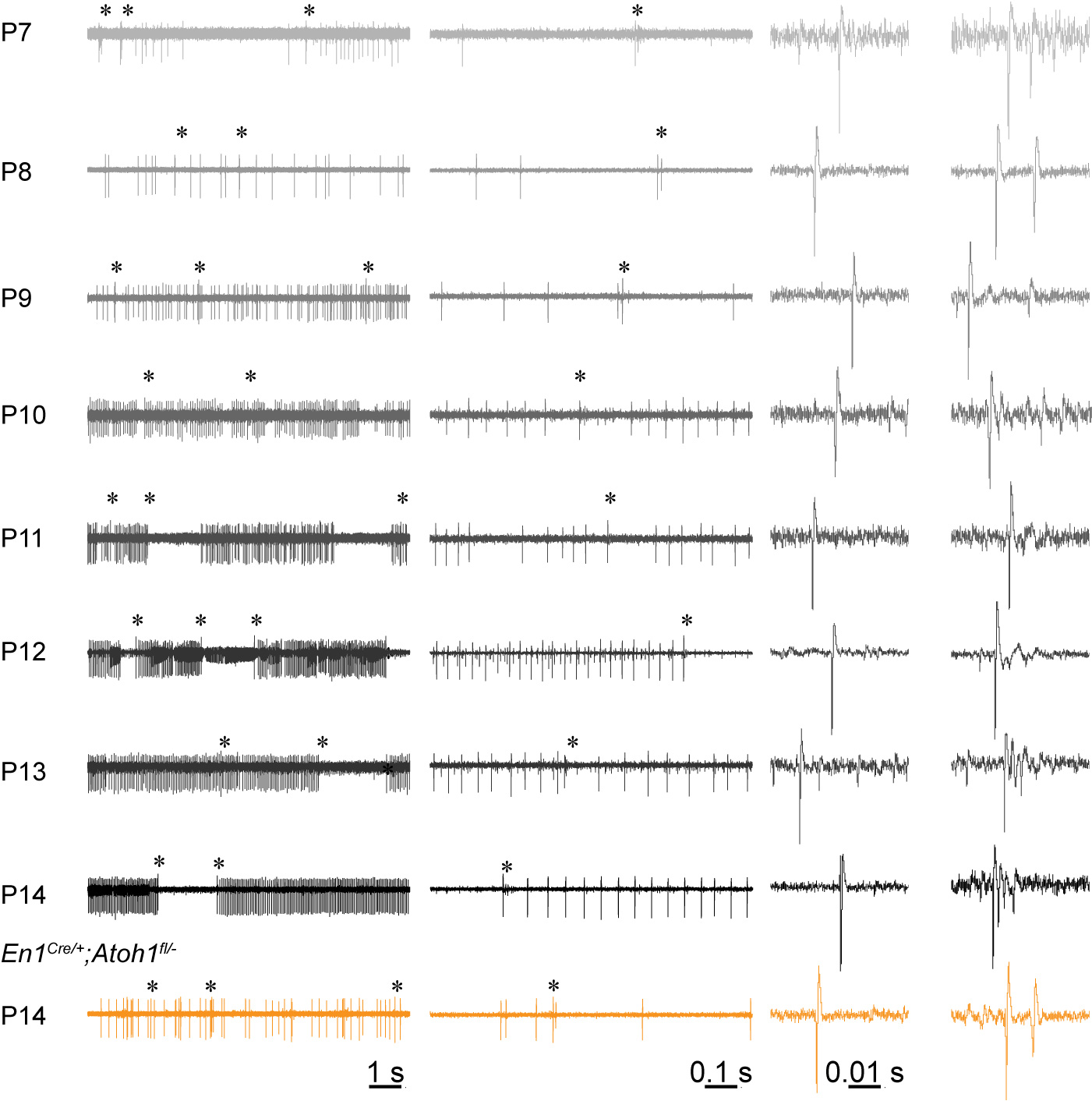
